# Altered network function in hippocampus after sub-chronic activation of Cannabis receptors in peri-adolescence

**DOI:** 10.1101/2025.09.16.676661

**Authors:** Johanna Rehn, Lucas Admeus, Bernat Kocsis

**Affiliations:** Dept Psychiatry at BIDMC, Harvard Medical School; Sahlgrenska Academy, University of Goteborg

## Abstract

The Cannabinoid 1-receptor (CB1R) is found in particularly high levels in the hippocampus (HPC), increased CB1R density and binding observed in patients with schizophrenia, and epidemiological studies suggest that regular cannabis use during adolescence is a risk factor for the disease. Historically, concerns around adolescent marijuana use focused on the development of psychosis later in life, however recent findings indicate that cognitive domains may also be at risk. CB1R was shown to interfere with neuronal network oscillations and to impair sensory gating and memory function. Neuronal oscillations are essential in multiple cognitive functions and their impairment was documented in neurological and psychiatric diseases. The aim of this study was to investigate how adolescent pre-treatment with the CB1R selective agonist CP-55940 may lead to abnormalities in theta synchronization in adulthood. Rats were pre-treated with CP-55940 (n=11, 6 males, 5 females) or vehicle (n=8, 4 males, 4 females) during adolescence (daily i/p injections between PND 32-36 or PND 42-46, n=10 and 9, respectively). They were then tested in adulthood (PND 70-88, n=17 or PND 111-115, n=2) under urethane anesthesia. Hippocampal theta rhythm was elicited by brainstem stimulation at 5 intensity levels one hour before and up to 5 hours after injection. We found a lasting significant decrease in theta power after CP-55940 in adult rats which was aggravated further in rats pre-treated in adolescence with the CB1R agonist. The effect was significantly larger (p=0.0462) in rats pre-treated during early adolescence (PND 32-36) compared to the group pre-treated during late adolescence (PND 42-46). We conclude that 1. Exposure to cannabis during adolescence leads to increased sensitivity to CB1R agonist in adulthood; 2. Early adolescence, a critical period for development of HPC networks generating theta rhythm, is particularly prone to this sensitivity.

## 1. Introduction

Historically, concerns around adolescent marijuana use focused on the development of psychosis later in life, however recent findings indicate that cognitive domains may also be at risk. The Cannabinoid 1-receptor (CB1R) is found in particularly high levels in forebrain structures, including the hippocampus (HPC) ^1^. Analysis of transcriptional regulatory factors of the endocannabinoid system has showed selective alterations of DNA-methylation of the gene CNR1, which codes for CB1R, in schizophrenic patients ^2^. This could indicate a potential upregulation of CB1R, an assumption that has been tested through several binding-analysis studies of cerebral CB1R ^3-5^.

On the other hand, CB1R was also shown to interfere with neuronal network oscillations and to impair sensory gating and memory function ^6-8^. Neuronal oscillations are essential in multiple cognitive functions and their impairment was documented in neurological and psychiatric diseases. There is a dense population of CB1R located on hippocampal GABAergic interneurons and upon presynaptic activation, CB1R inhibits the release of neurotransmitters, which leads to a modulation of GABA and Glutamate transmission ^9^, indicating that CB1R activation modulates the balance between inhibition and excitation of neural networks, which potentially causes desynchronization in the theta and gamma frequency bands ^10^.

Self-medication with cannabis is a recurrence in patients with schizophrenia, which lead to investigation of its potential to alleviate symptoms. Instead, it was found that cannabis exacerbates cognitive symptoms in patients with schizophrenia and can produce transient schizophrenic-like manifestations in healthy individuals ^11^. Epidemiological studies have also established that regular cannabis use during adolescence is a risk factor for the precipitation of the disease ^12^. The vulnerability of this developmental period is not limited to humans; studies using rats have shown significant changes in behavioral parameters following chronic peripubertal treatment with CB1R agonists which were not reproducible in adults ^13^. Adolescence represents a period where CB1R expression is at its peak and successively decrease up until adulthood in both rodents ^14^ and humans ^15^, which may indicate that during adolescence there is heightened susceptibility to exogenous ligands. Studying adolescent cannabinoid receptor-mediated modifications on neuronal functioning may thus provide useful insights into the mechanisms behind the pathophysiology of diseases such as schizophrenia.

The aim of this study was to investigate how adolescent pre-treatment with the CB1R selective agonist CP-55940 may lead to abnormal theta synchronization in adulthood. Two of our previous findings served as scientific premises of the project. First, CP-55940 significantly reduced theta and gamma power in HPC in adult freely behaving and theta in anesthetized rats, the latter reversed by CB1R antagonist AM-251 ^6^ and second, we have shown recently ^16^ in urethane anaesthetized rats that early adolescence is critical for development of HPC networks generating theta rhythm. We had reported that HPC theta power rapidly changes through postnatal days PND 32-39 with large interindividual variations, but stabilizes by late adolescence (PND 41-49) and then remains persistent until adulthood. Indeed, here we found massive aggravation of theta suppression induced by acute CP-55940 in adulthood if rats had been pre-treated with the CB1R agonist in adolescence. Furthermore, the effect was significantly larger in rats pre-treated during early adolescence (PND 32-36) compared to the group pre-treated during late adolescence (PND 42-46), i.e. matching the age, critical for development of oscillatory networks in the HPC.

## 2. Methods

All procedures were approved by the Institutional Animal Care and Use Committee (IACUC) of Beth Israel Deaconess Medical Center and conducted in accordance with the ARRIVE 2.0 guidelines. Twenty Sprague–Dawley rats (10 males, 10 females) were used (1 female was excluded in adult stage). Animals were housed under standard laboratory conditions with ad libitum access to food and water, maintained on a 12 h light/dark cycle. Body weights during the adolescent injection phase ranged from 95–198 g over the 5-day dosing period; at the time of electrophysiological recordings in adulthood, male rats weighed 375–500 g and females 125–420 g.

### 2.1. Drug preparation and administration

The selective CB1 receptor agonist CP-55940 (Sigma-Aldrich) was dissolved in 0.5% methylcellulose, which also served as vehicle for control injections. The working solution (0.3 mg/mL) was prepared by dissolving 10 mg CP-55940 in vehicle, followed by dilution to the target concentration. Rats received intraperitoneal injections of CP-55940 at a dose of 0.3 mg/kg in a volume of 1 mL/kg. Vehicle controls received methylcellulose alone in the same volume.

Animals were divided into four groups with balanced sex representation: early CP-55940 (n = 6), early control (n = 4), late CP-55940 (n = 5, 1 female was excluded at the stage), and late control (n = 4). Early administration (0.3 m/kg daily i/p injections)occurred on PND 32–36, and late administration on PND 42–46. The 5-day regimen was selected to model repeated adolescent cannabinoid exposure.

### 2.2. Anesthesia and surgical procedures

In adulthood, animals were anesthetized for electrophysiological recording using intraperitoneal urethane (2 g/kg total, given in two doses of 1 g/kg one hour apart). When required, a supplemental ketamine injection (0.05 mg/kg, i.p.) was administered to achieve a surgical plane of anesthesia. Ketamine was also used as the terminal agent before decapitation at the conclusion of experiments.

Following induction, the animals were secured in a stereotaxic frame. The skull was exposed and craniotomies were made above target regions. Screw electrodes were implanted for EEG recordings from cortical sites: frontal cortex, parietal cortex, a reference electrode anterior to the frontal site, and a ground electrode positioned over the cerebellum. For local field potential (LFP) recordings, stainless-steel fine-wire electrodes (125 µm) were stereotaxically implanted in the prefrontal cortex (PFC) and dorsal HPC according to Paxinos and Watson’s rat brain atlas. Twisted-pair electrodes were used to access both CA1 and dentate gyrus, allowing detection of phase reversals in hippocampal theta oscillations. A stimulating electrode was placed in the brainstem (reticular nucleus pontis oralis, RPO) using a guiding tube protruding ∼3 mm beyond the skull surface. Electrodes were fixed in place with dental acrylic, ensuring stable long-term recordings.

### 2.3. Electrophysiological recordings

Recordings were acquired with Dasylab 7.0 (Microstar Laboratories), converted to Spike2 (Cambridge Electronic Design), and stored for offline analysis. All channels were low-pass filtered at 20 Hz to remove high-frequency noise. Data were examined both in the time domain (waveform traces) and frequency domain (time–frequency spectrograms). Segments corresponding to RPO stimulation were extracted and subjected to Fast Fourier Transform (FFT) to compute power density spectra.

Signal amplitudes (in mV) can vary across subjects due to factors unrelated to neuronal activity, such as local bleeding, edema, or variability in electrode placement. To reduce this inter-animal variability, each dataset was normalized to its own pre-injection baseline activity. Theta power changes were assessed using paired t-tests comparing pre-versus post-injection recordings.

### 2.4. RPO stimulation to elicit HPC theta rhythm

RPO was stimulated by 0.1 ms square waves, 100 Hz trains of 10 s duration, separated by 1 min, using a programmable stimulator (Master 8, A.M.P.I.) through flexible isolator unit to generate constant current pulses (Iso-Flex, A.M.P.I.) First, the range of RPO-stimulation intensity was established in each particular experiment. These parameters (similar to LFP recordings in deep structures) depend on various technical factors, including the exact location of the stimulation electrodes, the condition of the ascending pathway, etc. and thus have to be set individually in each rat before the start of the experimental protocol. Two parameters were identified, the minimum (i.e. the threshold) stimulus eliciting theta rhythm in the HPC in most (>50%) trials, and the maximal stimulus beyond which no further increase in theta frequency is observed. The effective stimulus intensity varied between 0.04 and 1.0 mA in individual experiments; the range (i.e. max-min) was 0.57±0.04 mA. Thereafter repeated measurements of stimulation sequences at different intensity values within the two thresholds were performed. Each stimulation session consisted of 10 s-long, 1 per minute stimulation epochs, repeated 5 times at each of the 5 intensity levels, in random order, thus completing a stimulation session lasting ∼20 min). These sessions were executed in each rat in control condition immediately before the injection and then repeated once every hour for five hours after drug injection. All signals along with stimulation markers on a separate channel were recorded continuously during the entire experiment. For analysis, 8 s-long segments during stimulations were selected off-line, ignoring the potential transient induced by turning on and off the stimulator.

### 2.5. Statistical modeling

To evaluate treatment effects across animals, linear mixed-effects models were implemented in RStudio (Public Benefit Corporation). This approach was selected because mixed models handle missing values effectively, account for both fixed and random effects, and are less prone to inflated Type I error rates than repeated-measures ANOVA.

The primary model included pretreatment condition (CP-55940 vs. vehicle), time after injection, and dose as fixed effects. Random intercepts and slopes for individual animals were included to capture variability in baseline values and time-dependent responses. This specification assumes that animals vary both in baseline neural activity and in how their responses evolve over time, beyond what is explained by fixed effects.

Two additional models examined specific factors: one tested the effect of treatment timing (early vs. late adolescence), and another tested the effect of sex(male/female). Both models included time after injection as a fixed effect. Residuals were inspected visually with Q–Q plots and residuals-versus-fitted plots to confirm homoscedasticity and normality. Model selection was guided by Bayesian Information Criterion (BIC), which balances model fit against complexity. Lower BIC values were interpreted as evidence of superior model performance.

## 3. Results

To study the consequences of early life CB exposure on HPC oscillatory networks and to identify vulnerable periods of prepubertal neurodevelopment we inspected the changes in the acute reaction to CB1R activation in adult rats, previously subjected to sub-chronic CB exposure in adolescence (Fig.1B).

**Figure 1.**
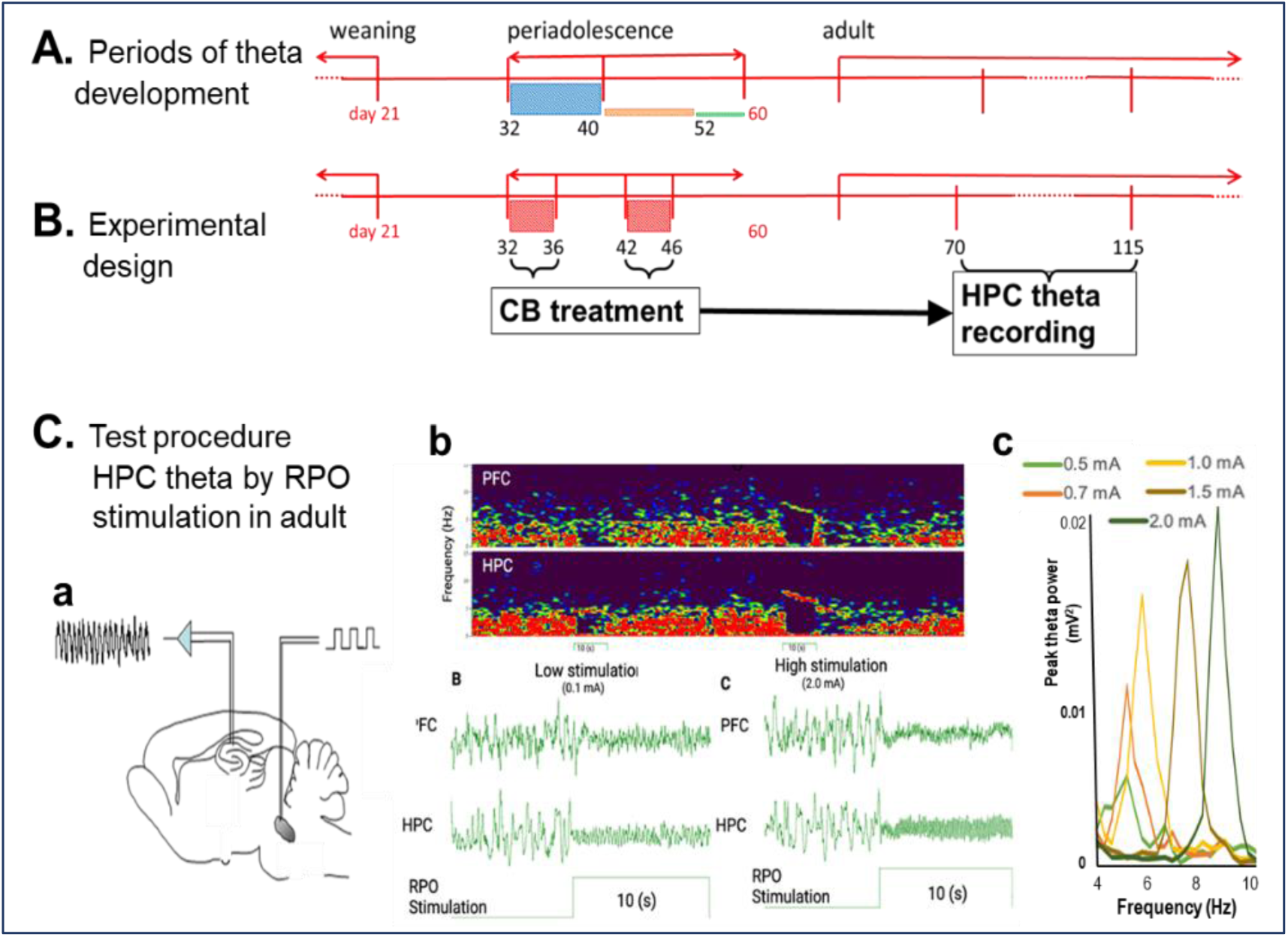
Experimental design (**A)**, based on critical periods of adolescent development of HPC oscillatory networks (**B**) of testing the consequences of CB exposure in adolescence on HPC theta reactions after the rats reached adulthood (**C**). **A-B**. CB exposure (xx mg/kg daily injections for 5 days) was performed either in PND 32-36 (n=10) when HPC theta activity is highly variable or in PND 42-46 (n=9), i.e. in the age when theta oscillations were shown established on a stable level (color columns through PND 32-40-52-60 in **A** are proportional to standard deviation of theta power within age groups^16^. **C**. Theta power was measured in adult rats (between PND 70-115), under urethane anesthesia, where HPC theta can be in induced in a controlled manner, i.e. independent of behavior, by electrical stimulation of the brainstem arousal pathways in the RPO (**a**) eliciting HPC theta rhythm (**b**) with increasing frequency (**c**) and power, depending on the stimulus intensity, individually identified in each experiment ^19^.

### 3.1. CB1R activation suppresses HPC theta oscillation in adult rats

It has been shown earlier, that acute administration of the selective CB1R agonist CP-55940 decreases HPC theta power in freely moving rats, correlated with memory impairment in HPC-dependent tasks^6, 8^. Suppression of spontaneous theta rhythm had also been reported in rats anesthetized with chloral hydrate^6^. Here we used urethane, which – of the anesthetics known not to restrain low-frequency (delta, theta) oscillations – has been most extensively used to investigate the neuropharmacology of networks generating theta rhythm^17-20^. In this model, HPC theta is elicited in a controlled manner by electrical stimulation of the brainstem arousal pathway to generate standard theta states, with progressive increase in theta frequency, amplitude, and power elicited with increasing stimulus intensity (Fig.1C, see Methods). Stable anesthesia under urethane lasts for hours and the reaction to brainstem stimulation remains unchanged. In this study, stimulation-induced theta was tested in ∼20 min segments every hour, before (see C1 in Figs.) and for five hours after drug injection (P1,…,P5). Lack of effect of vehicle injection stimulation-induced theta activity, widely reported in previous studies^17-20^, were verified in 2 naïve rats, untreated in peri-adolescence (Fig.2A).

**Figure 2.**
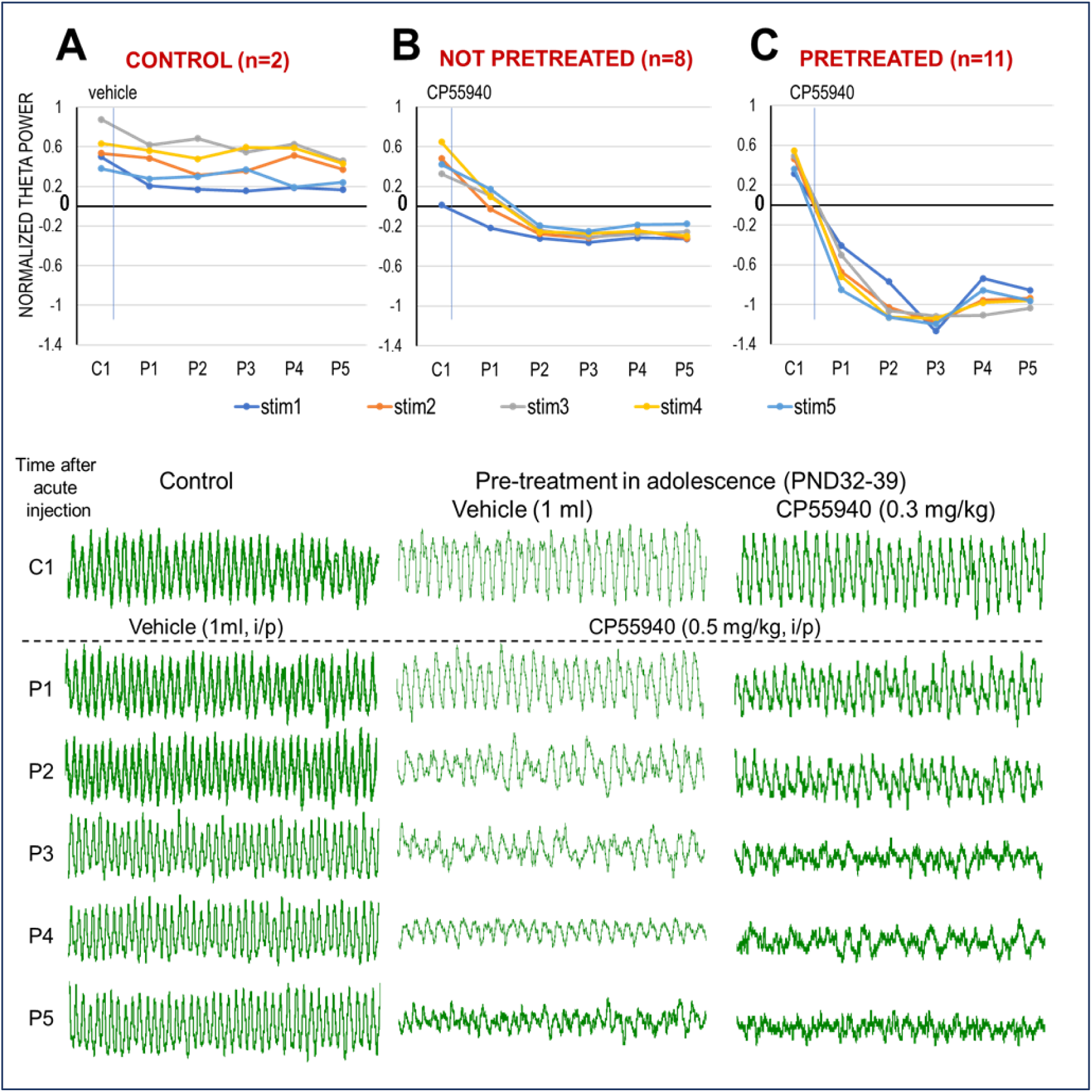
CB1R activation decreases HPC oscillation in adult rats. **A**. Stable, stimulus intensity-dependent (stim1…stim5) theta oscillations lasting for hours before (C1) and after vehicle injection (P1…P5). **B**. Effect of CB1R activation (0.5 mg/kg CP-55940 i/p injection) in adult rats with no prior CB exposure. Note reduction of theta power (group averages – *top panel*) and frequency (in the example – *bottom panel*) starting in P1 and more drastic from P2-P3. **C**. Effect of CB1R activation (0.5 mg/kg CP-55940) in adult rats injected with the same drug in adolescence (0.3 mg/kg, daily injections (*top*: PND 32-39 or 42-48 combined; *bottom*: sample treated in early adolescence). Note more severe reaction with no sign of theta rhythm in P3-P5. Theta power was normalized, setting the lowest and highest theta power, in each individual experiment, as 0 and 1, respectively. *Top panels*: group averages of theta power; *bottom*: examples of 10 s recordings during brainstem stimulation, in one experiment from each group.

First, the selective CB1R agonist CP-55490 was administered in 8 rats which had no prior exposure to the drug. HPC theta power started to decline by P1 (i.e. the first test, an hour after injection) and stabilized at a low level for the remaining of the recordings (P2 to P5, Fig.2B). The decrease in RPO-stimulated hippocampal theta power was highly significant (p < 0.05; Student’s paired t-test) for all stimulation intensity levels at all hourly tests, post-injection. Stimulation-elicited theta power was significantly below pre-injection control; it was consistently below even the smallest theta component recorded in C1 (set as 0 when normalizing the autospectra in individual experiments before group statistics).

The drastic main effect was slightly modulated by the dose of the CB1R agonist administered and by sex differences. CP-55940 was injected in concentrations of 0.3 mg/kg or 0.5 mg/kg (n=4 in each group). It was effective in both doses; theta power significantly decreased within an hour, starting at the first test, post-injection (P1), and lasted for at least 5 hours (P2 to P5). Although significant (p<0.05 at all stimulus levels, pairwise t-test), the effect of 0.3 mg/kg injection appeared less stable however; differences between the average theta power indicated a slower start (P1 in Fig.S1A) and a slight weakening of the effect 4-5 hours after injection (P4, P5). At the higher dose (0.5 mg/kg), CP-55940 drastically reduced theta power by P1 which then remained consistently low until the end of the experiments (Fig.S1B). Sex differences also manifested in a similar pattern of timing and stability of the reaction. Stimulation-elicited theta power was significantly below pre-injection controls at all stimulus intensities and all hourly tests in males as well as females (n=4 in each group). However, while in females, theta decreased right after the injection (P1) and stayed stable until the end (Fig.S1C), the reaction in males was smaller in P1 (yet, still significantly different from C1) and less stable 4-5 hours after injection (Fig.S1D).

### 3.2. Effect of pretreatment with CP-55940 in adolescence on HPC network oscillations

The decline in RPO-stimulated HPC theta power was considerably stronger after pretreatment with CP-55947 during adolescence (Fig.2C). Sample traces of the field potentials in three representative experiments, indicate that the acute effect of CP-55940 on theta rhythm was stronger in the adult rat exposed to cannabis in adolescence (0.3 ml/kg daily injections of CP-55940 for 5 days, PND 32-36 or PND 42-46). It displayed a reduction in theta amplitude in P1‒P2 and a total absence of visible theta rhythm three hours post-injection (Fig.2C) – in contrast with the progressive decline in theta amplitude in P1-P5 in the rat which had only received vehicle injections in adolescence (Fig.2B).

For statistical analysis of the effect of pretreatment with chronic administration of CB1R agonist during adolescence and the effect of various factors potentially modulating this effect, we used three linear mixed models for statistical group comparisons. This type of model is preferred over other tests (such as two-way ANOVA with replication) as it deals well with missing values and can include both fixed and random effects. Dividing the analysis into separate models decreases the risk of overfitting, while still allowing for exploration of the different sets of variables. The factor of primary interest, besides the main effect of adolescent exposure, were the timing of adolescent pretreatment to identify ages of vulnerability, and sex, to potentially reveal important details regarding the mechanism of the reaction.

In the primary model, the dynamics of post-injection theta power was tested with random intercept and random slope for time allowed for different subjects – with 3 variables, i.e. (1) pretreatment (ctrl vs. pretreated), the (2) testing dose (0.3 and 0.5mg/kg) and (3) time after acute injection (P1-P5) set as fixed effects. To justify the inclusion of a random intercept, it is assumed that the rats are allowed to have their own baseline values (and these are normally distributed around the average intercept). Regarding the random slope for time, we assume that the rats may respond to time differently. This could be due to inter-variability of how the rats may metabolize and respond to CP-55940, i.e. beyond what is explained by the fixed effects in the model. Overall, according to this model, the effect of adolescent exposure to cannabis, on adult theta reaction proved highly significant (p=0.022), as indicated by the differences in Figs.2B vs. 2C. Next, to test the effect of early (PND 32-36) and late PND 42-46) adolescent pretreatment as well as sex, two simpler mixed models were used with the fixed effect of time after injection as well as early/late pretreatment or male/female, respectively. Residuals were examined to confirm the assumption of constant variance, this was done by inspection of Q-Q plots as well as direct plots of the residuals against the fitted values. Furthermore, the Bayesian Information Criterion (BIC) was used to objectively compare the potential models based on their predictive capacity and complexity

We found that the age of sub-chronic CB1R activation in adolescence had a strong effect on the development of HPC networks (Fig.3). In both treatment groups, theta power was readily induced by RPO stimulation in adulthood (see e.g. C1 in Fig.3; also see traces in Fig.2C) and reacted to acute CP-55940 with fast decrease in power within an hour (P1) and did not recover for 5 hours. The levels of theta reduction were however different, depending on treatment age; the effect of early (PND 32-36) adolescent pretreatment with CP-55940 was significantly (p=0.0462, mixed model) stronger compared with the group pretreated during late (PND 42-46) adolescence (Fig.3A-B). Differences between not pretreated and early pretreated groups were also found significant (two-sample t-tests), at stimulus levels 2 through 5 (Stim4 shown in Fig.3C) in all post-stimulus trials (P1-P5). The effect of late adolescent exposure was considerably smaller and not well-elucidated by the less adequate t-tests; not significantly different from not-pretreated and occasionally not different from the early pretreated group either (e.g. in all trials at Stim3 and several trials at other stimulus levels; see e.g. in P1 and P4 at Stim4 shown in Fig.3C). The significant effects of the age of pre-treatment were detected by acute adult injection in adulthood using either the 0.3 or the 0.5 mg/kg dose.

**Figure 3.**
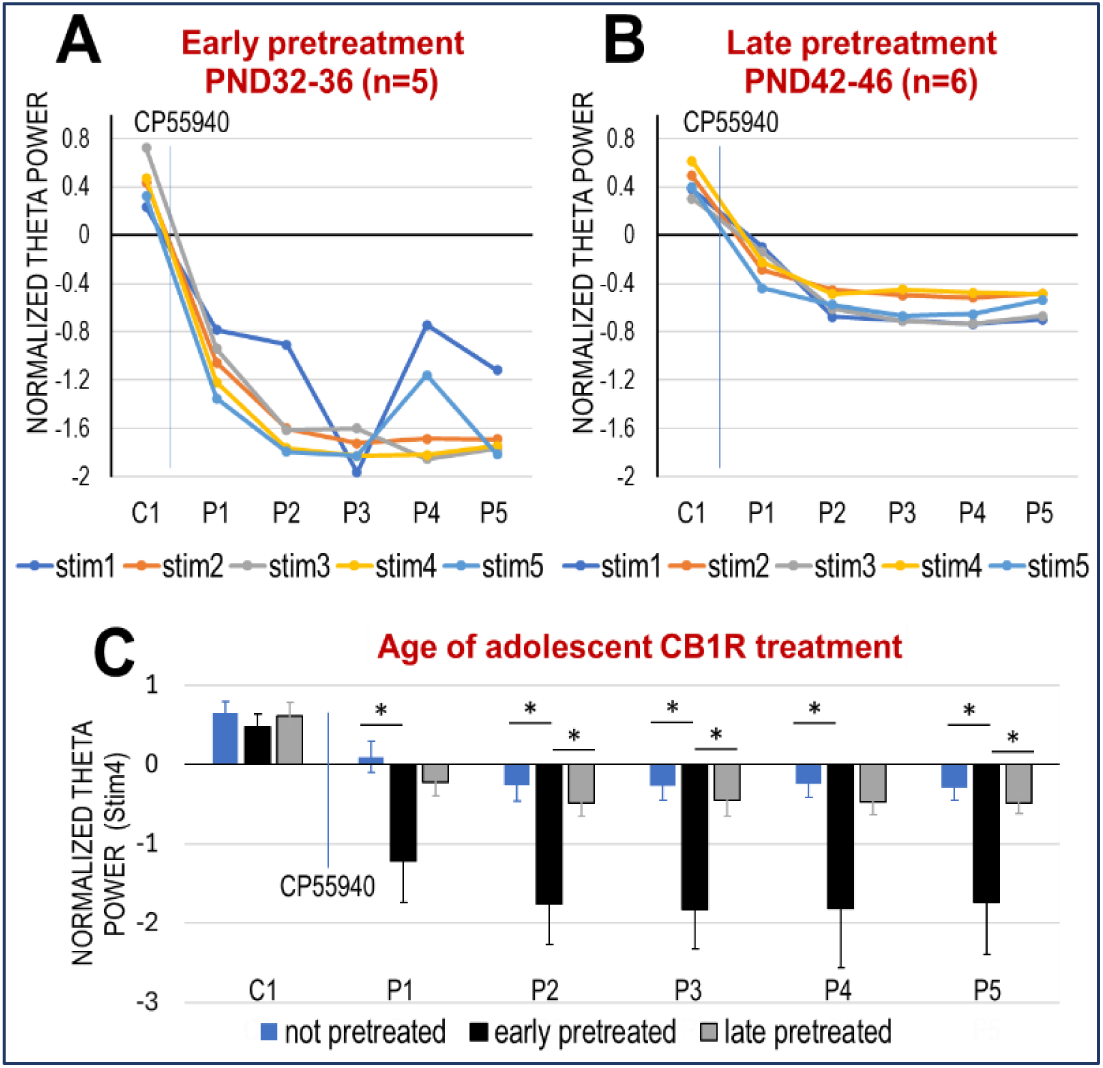
Effect of adolescent age of CB exposure on network oscillations in adulthood. **A-B**. Decrease of theta power (group averages) after acute CP-55940 injection in adult rats that had been exposed to the same drug (0.3 mg/kg, daily for 5 days) between PND 32-36 (**A**) or PND 42-46 (**B**). **C**. Comparison of changes in theta power elicited by stim4 after acute CP-55940 injection in CB naïve rats with rats exposed to CB1R agonist at different ages in adolescence (early-PND 32-36 or late-pretreated PND 42-46).

On the other hand, no difference was found in the reactions to CB1R activation between male and female rats pretreated with the same drug in adolescence (Fig.4A-B). Significant differences between the reactions to acute CP-55940 of pretreated and not pretreated rats was further verified by traditional two-sample t-tests in P1‒P5 at all stimulus levels (see e.g. Stim3, in Fig.4C)

**Figure 4.**
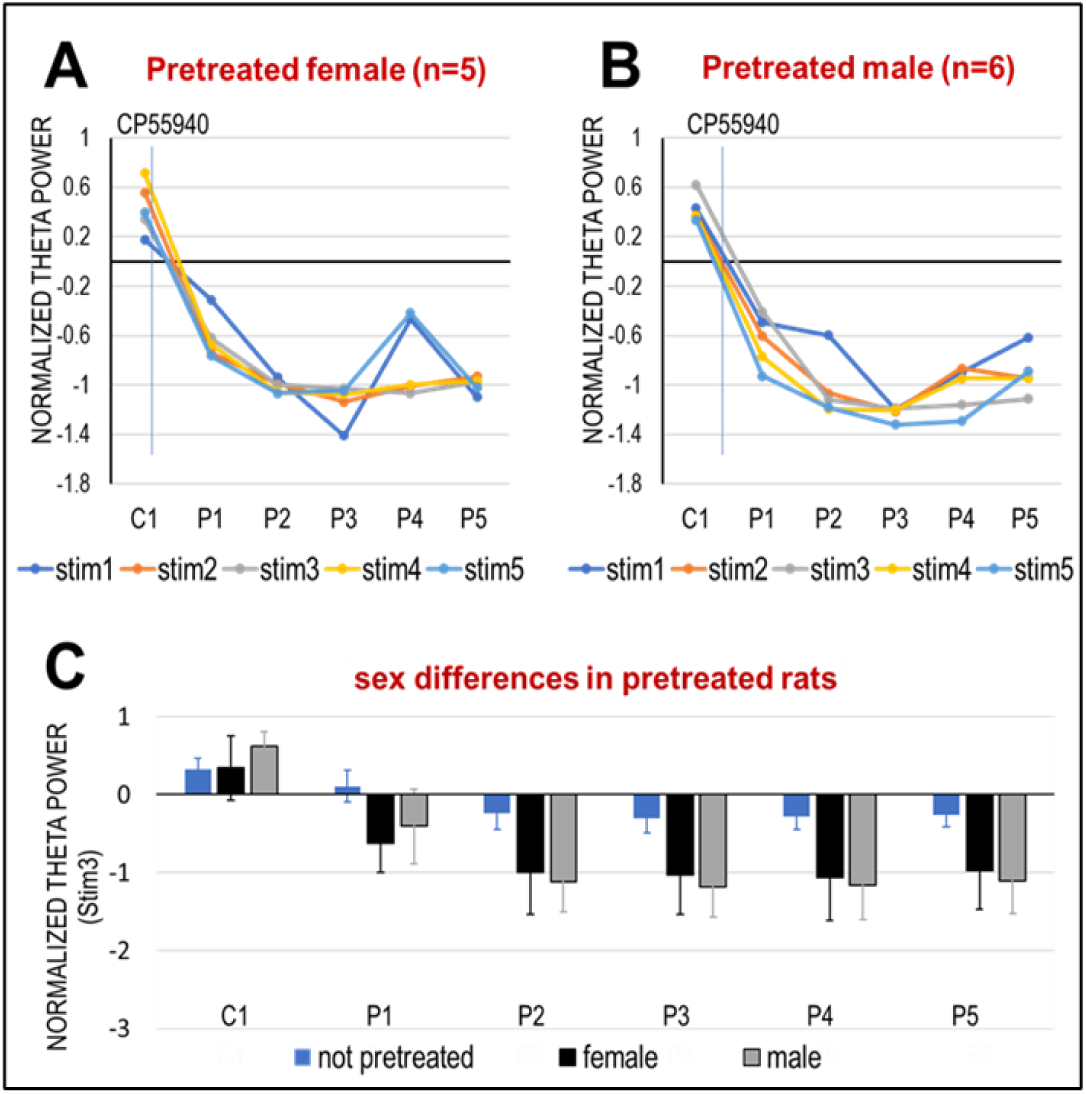
Effect of CP-55940 on elicited theta oscillations in female and male rats. **A-B**. Acute effect in rats pretreated in adolescence, group averages and SEM (**A**: females n=5, **B**: males n=6). **C**. Theta power elicited by Stim3 after acute injection of CP-55940 in adult rats pretreated with the same drug in adolescence (n=11).

## 4. Discussion

This study provides further evidence for the role of CB1R activation in the modulation of HPC network dynamics, demonstrating specifically heightened susceptibility of the theta generating networks in adulthood induced by sub-chronic adolescent exposure to CP-55940, thereby offering a potential neurobiological link to the epidemiological associations between early cannabis use and increased risk of developing schizophrenia. We found in particular, a lasting significant decrease in theta power induced by acute administration of CP-55940 in adult rats which was aggravated further in rats pre-treated in adolescence with the CB1R agonist. The effect was significantly larger in rats pre-treated during early adolescence (PND 32-36) compared to the group pre-treated during late adolescence (PND 42-46), i.e. matching the specific adolescent age critical for development of oscillatory networks in HPC ^16^.

The vulnerability of developmental periods in adolescence was demonstrated by changes in behavioral parameters following chronic peripubertal treatment of rats with CB1R agonists which were not reproducible in adults ^13^.

Although used in numerous prior studies, the technical protocols of sub-chronic CB administration in rodents during peri-adolescence have not been “standardized”. Adolescence in rats is typically defined as the entire period between PND 21 (weaning) to PND 60 (adulthood), with a subdivision of early (prepubescence, PND 21-34), middle (peri-pubescence, PND 34-46), and late (young adult, PND 46-59) adolescence ^22^. Prior studies testing the consequences of CB1R agonists exposure differed in the compound used (mostly CP-55940 ^23-26^ but also WIN-55212-2 ^13^ or THC ^27^), in the dose (0.3-0.7 mg/kg of CP-55940), and the timing (early ^25, 28^ and late ^13, 26^ adolescence) and length (3-10 ^25-30^ or >20 days ^13, 23, 24^) of daily injections. In this study, we used the highly selective agonist CP-55940 which is 45 times more potent than Δ^9^-THC, the CB1R agonist commonly used in research ^31^. As for the age of sub-chronic cannabis exposure, our choice was guided by the recent report by Sibilska et al.^16^, directly monitoring the developmental alterations in HPC theta oscillations during adolescence. Performed under urethane anesthesia, it followed a “pseudo-longitudinal” design using rats-siblings from the same mother, to reduce individual innate differences between subjects. Collecting HPC theta power measurements in these rats in each day of adolescence from PND 32 to PND 52, with progressively increasing body weights (from 115 to 190 grams in females and 120 to 310 grams in males), clearly demonstrated that stable theta rhythm was established by PND 40, after a period of progressive development in PNS 32-40 with high inter-subject variability. The correlation between age and theta power was significant in both males and females and theta stabilization was also apparent in both sex groups, but while in females it was complete by PND 40, in males this process was somewhat slower or delayed ^16^. The results of the present study revealed an augmented vulnerability of this early stage of development, further extending the overall effect of cannabis exposure over peri-adolescence.

Sex differences in maturation of theta-generating HPC networks ^16^ might also explain, at least in part, the lack of sex-related discrepancies in the current study, apparently contradictive to existing literature suggesting gender-specific differences in CB1R affinity and receptor distribution ^32^. As early adolescence (PND 32-36) is associated with a significantly higher susceptibility compared to later in adolescence (PND 42-46), if the critical period for stabilization of important theta-generating networks occurs before puberty ^33^, then the similarity between the sexes may seem less surprising. Traditionally, preferential use of male rodents has been pointing to uncertainties due to the variability induced by the estrous cycle. As a result, there is significantly more data available on male rodents; there is a strong bias towards excluding female subjects. Due to the heterogeneous nature of both schizophrenia ^34^ and cannabis-use ^35^, it is of great importance to test potential sex differences in future studies.

Although cannabis exposure in human is primarily associated with smoking marijuana, animal studies use daily injections which allow introducing pharmacological compounds with selective receptor action, and reliable drug delivery and dose control. This study also used minimally invasive intraperitoneal injections daily for 5 days, with no further manipulations; the pups receiving drug or vehicle were not even separated, were were maintained in common cages.

The testing procedure in adult rats however was performed under urethane anesthesia which although imposing certain constraints on the interpretation has well-defined advantages on pharmacological investigation of neuronal and network mechanisms, especially when the pharmacological manipulations (both adolescent treatment and acute test injection in adults, in our experiments) might induce changes in behavior not necessarily or directly linked with the specific neuronal level alterations under investigations. In freely moving animals, HPC theta rhythm is mainly associated with exploratory behavior and gross movements but may also occur in awake immobility, elicited by a range of sensory inputs which are hard to account for; in their classic paper for example, Kramis et al.^36^ reported theta activity in reaction even to “changes in the experimenter’s facial expression”. Thus, by effectively eliminating behavioral confounds of waking on one hand and fragmentation of sleep patterns (e.g. short REM sleep, never lasting longer than a few minutes in rodents) on the other, the major advantages of this preparation go beyond the “convenience” provided by an immobile, stable preparation.

The rationale for using the model of urethane-anesthetized rats in this study was that 1. unlike many other anesthetics, urethane leaves slow oscillations intact; 2. this model has been used for decades to study the mechanism of generation and control of theta rhythm, including drug studies with firmly established predictive validity (e.g. ^6, 20, 37-41^); 3. theta rhythm can be elicited by electrical stimulation of the ascending arousal pathway in a quantitively controlled manner, i.e. without behavioral confounds. 4. Yet, theta reaction to acute CB1R agonist in adult rodents was only tested earlier in mice under Chloral Hydrate anesthesia and thus the design of the present study included the full investigation of untreated normal adult rats to provide the necessary control. We found that acute CP-55940 did suppress theta rhythm in naïve rats in this model providing such a baseline to quantify the effect of adolescent cannabis exposure. RPO stimulation with increasing intensity is delivered in a precisely controlled manner, to mimic the action of ascending brainstem arousal pathways shown to have a role in maintaining rhythmic synchronized forebrain activity, such as HPC theta, specific to awake exploratory behavior and REM sleep.

Oscillatory synchronization is evolutionarily well preserved; all oscillations, including HPC theta, are present in the same frequency bands in mammals, including rodents and humans ^42^. Disrupted neuronal synchronization and coupling are markers of dysfunctional information processing in patients with schizophrenia and could help to explain the impaired cognitive functions observed in patients ^43^. Traditionally, schizophrenia research has focused on high frequency oscillations such as gamma ^44-46^ in active waking and more recently, on sleep spindles ^21, 47-49^, while the impairment of low frequency synchronization remains a highly understudied area. The impairment of theta oscillations during novelty-related tasks has been observed in schizophrenic patients ^50^ and disruption of the coupling between cortical areas and HPC is thought to arbitrate some of the pathophysiology in schizophrenia and finding mechanisms for this occurrence is of great importance ^51^.

## Supplemental Figure

**Figure S1.**
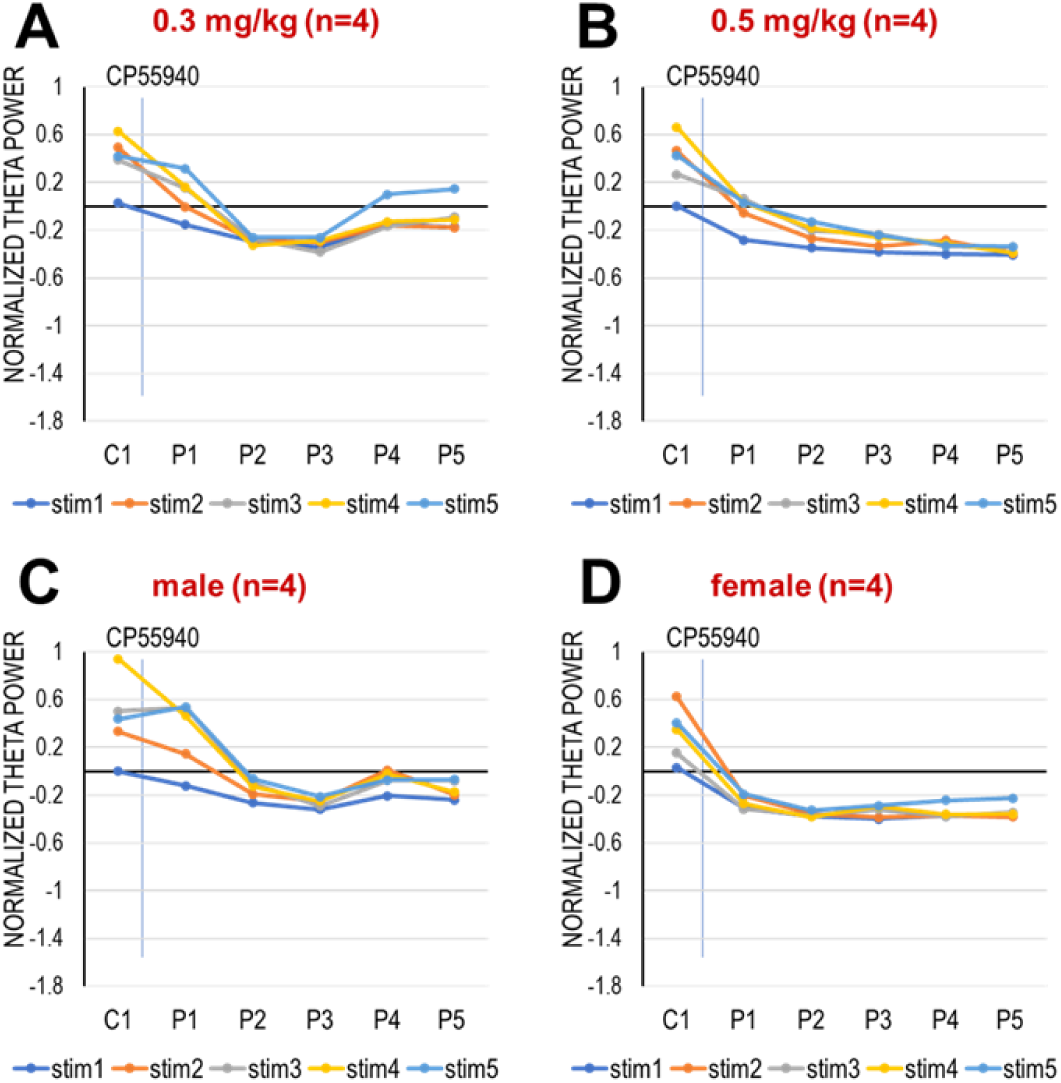
Effect of the testing dose (A-B) and sex (C-D) on the effect of CB1R activation in adult rats with no prior CB exposure. A-B. Effect of CP-55940 injected in different doses (0.3 and 0.5 mg/kg, A and B, respectively). C-D. Effect of CP-55940 injection in adult male (C) and female rats (D). Note slight differences of theta suppression only expressed in delayed onset (P4-P5) and less stable, shorter reaction (P2-P3) following the lower CP-55940 dose and in male rats.

## Acknowledgements

This work was supported by the National Institute of Mental Health grant R01 MH100820 to BK and a Swedish Government stipend to RM.

## Author contributions

RM and BK contributed to the experiments, analysis, interpreting the results, and writing the report; LA contributed to the analysis.

